# Global biochemical and structural analysis of the type IV pilus from the Gram-positive bacterium *Streptococcus sanguinis*

**DOI:** 10.1101/459388

**Authors:** Jamie-Lee Berry, Ishwori Gurung, Jan Haug Anonsen, Ingrid Spielman, Elliot Harper, Alexander M. J. Hall, Vivianne J. Goosens, Claire Raynaud, Michael Koomey, Nicolas Biais, Steve Matthews, Vladimir Pelicic

**Affiliations:** From the Medical Research Council Centre for Molecular Bacteriology and Infection, Imperial College London, London, United Kingdom; Department of Biological Sciences, Proteomics and Mass Spectrometry Unit, University of Oslo, Oslo, Norway; Department of Biological Sciences, Center for Integrative Microbial Evolution, University of Oslo, Norway; Department of Biology, Brooklyn College of the City University of New York, New York, USA; The Graduate Center of the City University of New York, New York, USA; Centre for Structural Biology, Imperial College London, London, United Kingdom

**Keywords:** Gram-positive bacteria, *Streptococcus sanguinis*, type IV filaments, type IV pili, pilin, protein structure, twitching motility, virulence factor, molecular motor

## Abstract

Type IV pili (Tfp) are functionally versatile filaments, widespread in prokaryotes, that belong to a large class of filamentous nanomachines known as type IV filaments (Tff). Although Tfp have been extensively studied in several Gram-negative pathogens where they function as key virulence factors, many aspects of their biology remain poorly understood. Here, we performed a global biochemical and structural analysis of Tfp in a recently emerged Gram-positive model, *Streptococcus sanguinis*. In particular, we focused on the five pilins and pilin-like proteins involved in Tfp biology in *S. sanguinis*. We found that the two major pilins, PilE1 and PilE2, (i) follow widely conserved principles for processing by the prepilin peptidase PilD and for assembly into filaments; (ii) display only one of the post-translational modifications frequently found in pilins, *i.e*. a methylated N-terminus; (iii) are found in the same hetero-polymeric filaments; and (iv) are not functionally equivalent. The 3D structure of PilE1, solved by NMR, revealed a classical pilin fold with a highly unusual flexible C-terminus. Intriguingly, PilE1 more closely resembles pseudopilins forming shorter Tff than *bona fide* Tfp-forming major pilins, underlining the evolutionary relatedness among different Tff. Finally, we show that *S. sanguinis* Tfp contain a low abundance of three additional proteins processed by PilD, the minor pilins PilA, PilB, and PilC. These findings provide the first global biochemical and structural picture of a Gram-positive Tfp and have fundamental implications for our understanding of a widespread class of filamentous nanomachines.

---

Type IV pili (Tfp) are thin, long and flexible surface-exposed filaments, widespread in Bacteria and Archaea, which mediate a wide array of functions (1,2). Tfp are polymers of primarily one major protein subunit - a type IV pilin with distinctive N-terminal sequence motif (class III signal peptide) and 3D structure (3) - assembled by a conserved set of dedicated proteins (4). These defining features are shared by a large class of filamentous nanomachines called type IV filaments (Tff) (1). Tff are ubiquitous in prokaryotes since genes encoding type IV pilins and filament assembly proteins are found in virtually every bacterial and archaeal genome (1).

Much of our current understanding of Tff biology comes from studies of bacterial Tfp in a few Gram-negative human pathogens in which they function as key virulence factors (4). The following general picture has emerged. Prepilins are translocated by the general secretory pathway across the cytoplasmic membrane (5,6), where they remain embedded via a universally conserved structural feature, *i. e*. a protruding hydrophobic N-terminal α-helix (α1N) (3). This leaves the hydrophylic leader peptide of the class III signal peptide in the cytoplasm, which is then cleaved by a membrane-bound aspartyl protease (7) - the prepilin peptidase - generating a pool of mature pilins in the membrane ready for polymerisation into filaments. Efficient prepilin processing by the prepilin peptidase, which does not require any other protein (8), depends on the last residue of the prepilin leader peptide (a conserved Gly) (9) and two conserved catalytic Asp residues in the prepilin peptidase (7,10). How filaments are polymerised remains poorly understood, but it is clear that this process is mediated by a multi-protein machinery in the cytoplasmic membrane (11,12), which transmits energy generated by a cytoplasmic hexameric assembly ATPase (13) to membrane-localised pilins. As a result, pilins are extruded from the membrane and polymerised into helical filaments via hydrophobic packing of their α1N helix within the filament core (14,15). Finally, once Tfp reach the outer membrane, they are extruded onto the surface through a multimeric pore - the secretin (12,16,17). The above picture, although complex, is oversimplified because there are additional proteins that play key roles in Tfp biology, including several proteins with class III signal peptides, named minor pilins or pilin-like proteins, whose localisation and exact role are often unclear. Moreover, Tfp are highly dynamic filaments, constantly extending and retracting. Retraction has been best characterised in a sub-class known as Tfpa, where it results from filament depolymerisation powered by the cytoplasmic hexameric ATPase PilT (18), which generates massive tensile forces (19,20).

Until recently, Tfp have not been extensively studied in Gram-positive species although it has been recognised that this represents a promising new research avenue as these bacteria possess a simpler surface architecture (21). Although Gram-positive Tfp were first described in Clostridiales (22,23), *Streptococcus sanguinis* has emerged as a model because it is genetically tractable (24,25). A comprehensive genetic analysis of *S. sanguinis* Tfpa (24) has revealed that they: (i) are assembled by a similar machinery as in Gram-negative species but with fewer components, (ii) are retracted by a PilT-dependent mechanism, generating tensile forces very similar to those measured in Gram-negative species, and (iii) power intense twitching motility. The main peculiarity of *S. sanguinis* filaments is that they contain two major pilins, rather than one as normally seen (24).

In the present study, we have focused on the pilins and pilin-like proteins involved in Tfp biology in *S. sanguinis* and have performed a global biochemical and structural analysis of its filaments.

## Results

### The pil locus in S. sanguinis 2908 encodes five pilin/pilin-like proteins

All known integral components of Tfp and/or Tff share an N-terminal sequence motif named class III signal peptide (1,3). It consists of a leader peptide, composed predominantly of hydrophilic amino acids (aa) ending with a conserved Gly_-1_, followed by a stretch of 21 predominantly hydrophobic aa (except for a negatively charged Glu_5_) forming the protruding α1N helix that is the main assembly interface for subunits within filaments (1,3). Processing by the prepilin peptidase PilD occurs after Gly_-1_. In most Tfp and/or Tff, there are mutiple pilin and/or pilin-like proteins (1,3). Bioinformatic analysis of the proteins encoded by the *pil* locus in *S. sanguinis*, which contains all the genes involved in Tfp biology (24), predicts five pilins and/or pilin-like proteins (PilA, PilB, PilC, PilE1 and PilE2) (Fig. S1A). Four proteins (PilB, PilC, PilE1 and PilE2) display a canonical N-terminal IPR012902 motif (26), which is part of the class III signal peptide (Fig. 1A). Visual inspection of the sequences of the remaining proteins suggests that PilA has a degenerate class III signal peptide (Fig. 1B), which is not identified by the bioinformatic tools available, including PilFind that is dedicated to the identification of type IV pilins (27).

**Figure 1.**
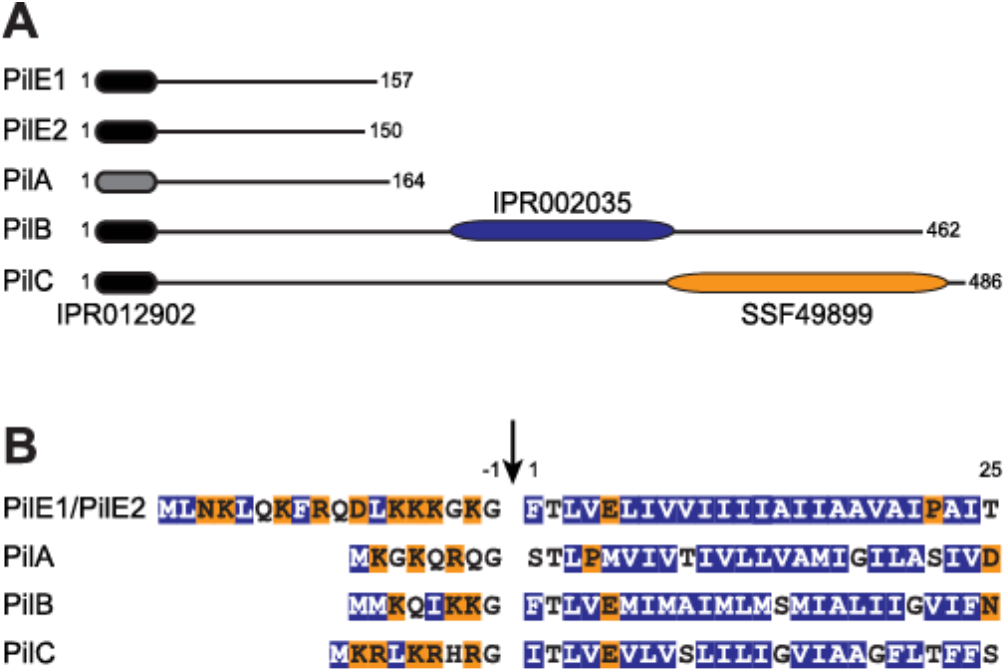
Bioinformatic analysis of the two major pilins (PilE1 and PilE2) and three pilin-like proteins (PilA, PilB and PilC) encoded by the *pil* locus in *S. sanguinis* 2908. (**A**) Protein architecture of the Pil proteins harbouring the N-terminal IPR012902 motif that is part of the class III signal peptide defining type IV pilins (black rounded rectangle). In PilA, this motif could be detected only by visual inspection and is therefore represented by a grey rounded rectangle. PilB and PilC contain additional C-terminal domains (blue and orange rounded rectangles). PilB contains a von Willebrand factor type A motif (IPR002035), while PilC contains a concanavalin A-like lectin/glucanase structural domain (SSF49899). Proteins have been drawn to scale and the subscript numbers indicate their lengths. (**B**) N-terminal sequence alignment of the putative class III signal peptides of the above five Pil proteins. The 8-18 aa-long leader peptides, which contain a majority of hydrophilic (shaded in orange) or neutral (no shading) residues, end with a conserved Gly_-1_. Processing by PilD (indicated by the vertical arrow) is expected to occur after Gly_-1_. The mature proteins start with a tract of 21 predominantly hydrophobic residues (shaded in blue), which usually forms an extended α-helix that is the main assembly interface within filaments.

PilA, PilB and PilC - all of which are essential for piliation (24) - exhibit sequence features distinct from PilE1 and PilE2, the major pilin subunits of *S. sanguinis* Tfp (24). All three proteins have much shorter leader peptides than PilE1/PilE2 (Fig. 1B). In addition, although mature PilA has a size (17 kDa) similar to previously studied pilins and pilin-like proteins (3), its putative class III signal peptide is unique for several reasons. The first residue of the predicted mature protein is a Ser_1_, there is an unusual Pro4, and the highly conserved Glu_5_ is missing. On the other hand, while PilB and PilC have canonical class III signal peptides (Fig. 1B), both are much larger than classical pilin-like proteins, with mature sizes of 50 and 52.7 kDa, respectively (Fig. 1A). This is explained by the presence of bulky domains at the C-terminus of PilB and PilC. PilB contains a von Willebrand factor type A motif (IPR002035) (26), while PilC contains a concanavalin A-like lectin/glucanase structural domain (SSF49899) (28) (Fig. 1A).

Taken together, these findings suggest that PilA, PilB and PilC are pilin-like proteins, which might be cleaved by PilD and polymerised into *S. sanguinis* Tfp, alongside the major pilins PilE1 and PilE2.

### S. sanguinis Tfp are hetero-polymers of N-terminally methylated PilE1 and PilE2

As shown previously, purified *S. sanguinis* Tfp consist predominantly of comparable amounts of two proteins, the major pilins PilE1 and PilE2 (24) which share extensive sequence identity (Fig. S2). Tfp, sheared by vortexing, were purified by removing cells/cellular debris by centrifugation before pelleting filaments by ultra-centrifugation (24). As assessed by Coomassie staining after SDS-PAGE (Fig. 2A), these pilus preparations contain contaminants that originate from cells/cellular debris. In order to determine the precise composition of *S. sanguinis* Tfp, we improved the purity of our pilus preparations. To remove more debris, we performed one additional centrifugation step and passed the sheared filaments through a 0.22 μm syringe filter before ultra-centrifugation. As assessed by SDS-PAGE/Coomassie (Fig. 2A), filaments prepared using this enhanced purification procedure were significantly purer, with no visible contaminants. Intriguingly, the morphology of the purified filaments also changed. While they were previously overwhelmingly thick (12 nm) and wavy (24), most filaments purified using this enhanced procedure display a classical Tfp morphology (1,4) as assessed by transmission electron microscopy (TEM) (Fig. 2B). Indeed, they are thin (~6 nm wide), long (several μm) and flexible, but they do not form large bundles.

**Figure 2.**
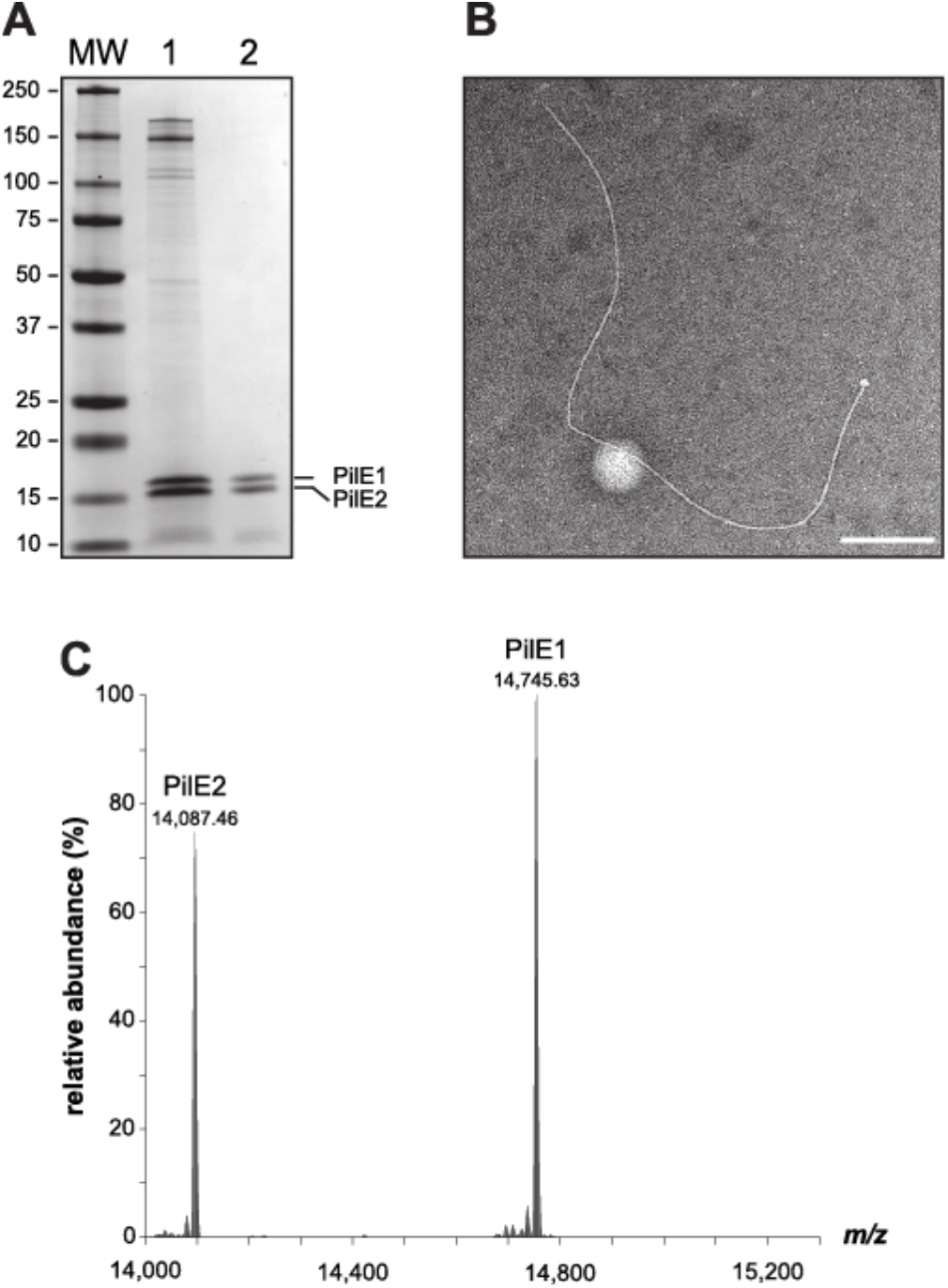
*S. sanguinis* Tfp exhibit a characteristic Tfp morphology and are composed primarily of N-terminally methylated PilE1 and PilE2 subunits. (**A**) SDS-PAGE/Coomassie analysis of *S. sanguinis* Tfp purified using previously described (lane 1) and enhanced (lane 2) purification procedures. Samples were prepared from cultures at similar OD_600_, separated by SDS-PAGE and stained with Coomassie blue. Identical volumes were loaded in each lane. MW indicates the molecular weight marker lane, with weights in kDa. (**B**) Filament morphology in WT pilus preparations prepared using the enhanced purification procedure as assessed by TEM after negative staining. Scale bar represents 200 nm. (**C**) Top-down mass profiling of purified Tfp from *S. sanguinis*. Shown are deconvoluted molecular mass spectra over the range of 14,000 to 15,300 *m/z*. Masses are presented as monoisotopic [M+H]^+^.

In other piliated species, major pilins frequently undergo post-translational modifications (PTM) (3). Their N-terminal residue upon processing is usually methylated and other PTM sometimes include the addition of a variety of glycans and phosphoforms. Methylation is catalysed by PilD, which is a bi-functional enzyme in many species (8,9). A bioinformatic analysis shows that *S. sanguinis* PilD contains the IPR010627 motif catalysing N-methylation (Fig. S1B), suggesting that the N-terminal Phe1 residue in PilE1 and PilE2 is likely to be methylated. Since other PTM cannot be inferred bioinformatically, we used top-down and bottom-up mass spectrometry (MS) to map the PTM of the two major pilins of *S. sanguinis* 2908. Top-down MS analysis of purified filaments showed the presence of two major proteoforms with a ratio of approximately 4:3 (PilE1:PilE2) (Fig. 2C), with deconvoluted singly charged monoisotopic masses at *m/z* 14,745.63 and 14,087.46 Da. These masses are consistent with the predicted theoretical masses of mature PilE1 (14,731.60 Da [M+H]^+^) and PilE2 (14,073.40 Da [M+H]^+^) with the addition of a single N-terminal methyl group (14.01 Da). Bottom-up LC-MS/MS analysis of the two bands excised separately and digested in-gel identified PilE1 and PilE2 with nearly complete sequence coverage (85% and 86%, respectively). The only peptide that was not detected, despite employing several proteolytic enzymes, was the N-terminus (_1_FTLVELIVVIIIIAIIAAVAI_21_) (Fig. S2). These MS results strongly suggest that *S. sanguinis* pili consist mainly of a 4:3 ratio of N-terminally methylated PilE1 and PilE2 subunits.

Next, we sought to answer the question whether *S. sanguinis* Tfp are heteropolymers of PilE1 and PilE2 or whether two homo-polymers co-exist. Recently, using a markerless gene editing strategy (25), we showed that *pilE1* could be engineered *in situ* to encode a protein with a C-terminal 6His tag, without affecting piliation or Tfp functionality. We therefore used this property to design an affinity copurification procedure to answer the above question (Fig. 3A). In brief, the intention was to (i) engineer *pilE1* and *pilE2* mutants encoding C-terminally 6His-tagged proteins, (ii) shear the filaments, (iii) affinity-purify (pull down) sheared filaments containing 6His-tagged subunits, and (iv) assess whether the untagged pilin co-purifies, suggesting that the filaments are hetero-polymers, or does not co-purify, suggesting that two distinct homopolymers co-exist (Fig. 3A). We therefore engineered four different variants by either fusing a 6His tag to the C-terminus of full-length PilE1 and PilE2 (PilE1_6His-long_ and PilE2_6His-long_) or by replacing the last seven aa in these pilins by the tag (PilE1_6His-short_ and PilE2_6His-short_). We first confirmed by SDS-PAGE/Coomassie analysis of purified pilus preparations that the four variants were piliated (Fig. 3B), although at various levels. Purified pili contained both tagged and untagged pilins as assessed by immunoblotting using antibodies specific for PilE1 and PilE2 (24), or an anti-6His tag commercial antibody (Fig. 3B). When affinity-purified sheared filaments were analysed by immunoblotting using anti-PilE1, anti-PilE2 or anti-6His antibodies, we found that in each of the four mutants the untagged pilin co-purifies with the tagged pilin (Fig. 3C). Importantly, wild-type (WT) untagged filaments cannot be affinity-purified using this procedure, indicating that the pull down is a 6His tag-specific process (Fig. 3C). Considering that *S. sanguinis* Tfp show little if any bundling, these findings strongly suggest that they are hetero-polymers of PilE1 and PilE2.

**Figure 3.**
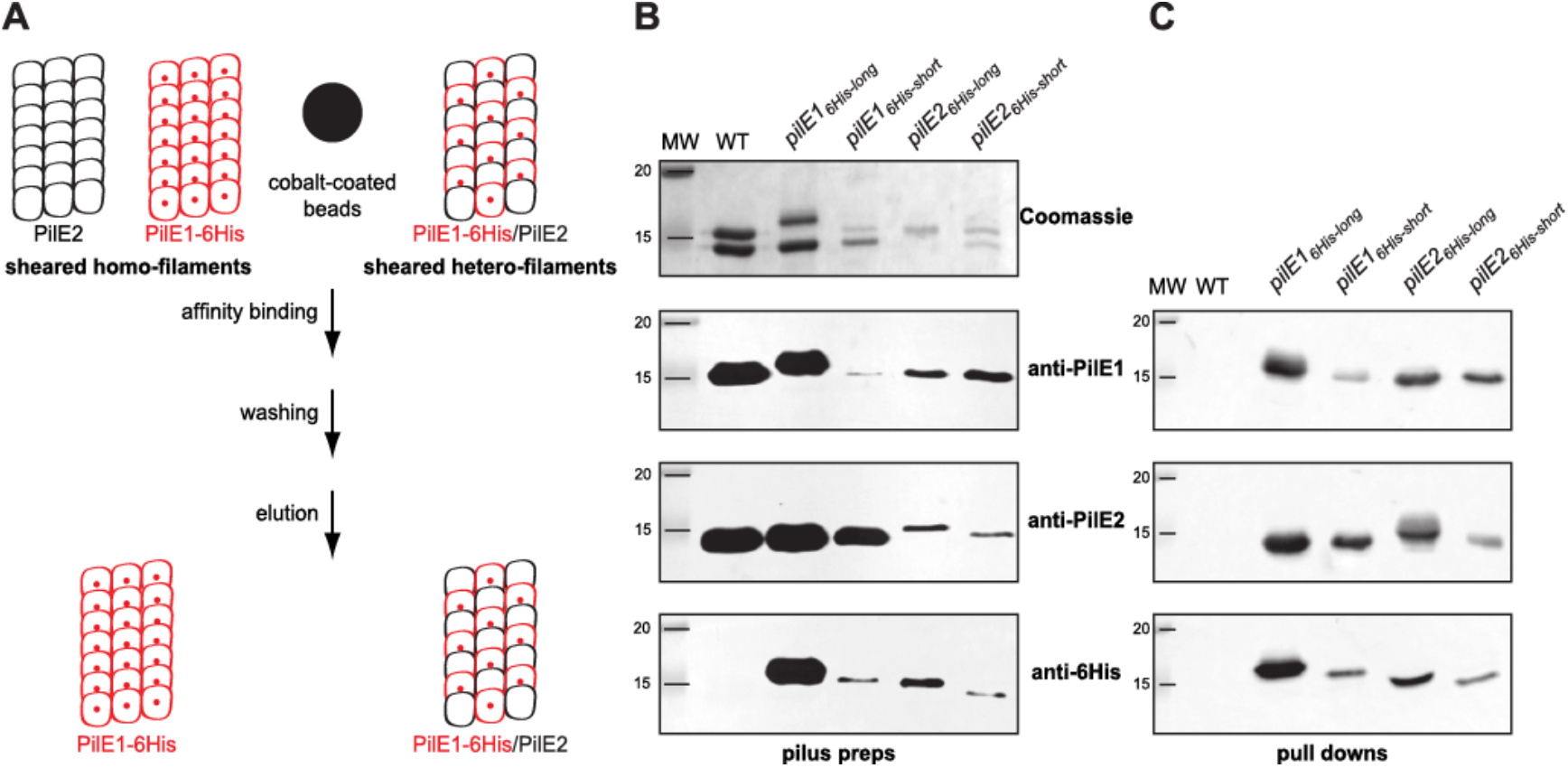
*S. sanguinis* Tfp are hetero-polymers composed of PilE1 and PilE2. (**A**) Schematics of the affinity-purification strategy. In brief, one of the genes encoding major pilin PilE1 (red) or PilE2 (black) is engineered to produce a protein fused to an affinity 6His tag, *e.g* PilE1-6His (small red sphere). Sheared pili are mixed with cobalt-coated beads (large black sphere) and purified by pull down, which is expected to yield filaments containing both pilins (tagged and untagged) if *S. sanguinis* Tfp are hetero-polymers. Conversely, if the filaments are distinct homo-polymers, only the tagged filaments will be purified. (**B**) Analysis of filaments purified by shearing/ultra-centrifugation from four unmarked mutants harbouring a 6His tag fused either to the C-terminus of full-length PilE1 and PilE2 (PilE1_6His-long_ and PilE2_6His-long_) or replacing the last seven aa in these pilins (PilE1_6His-short_ and PilE2_6His-short_). The WT strain was included as a control. Samples were prepared from cultures at similar OD_600_, separated by SDS-PAGE and either stained with Coomassie blue (upper panel) or analysed by immunoblot using anti-PiE1, anti-PilE2 or anti-6His antibodies (bottom three panels). MW indicates the molecular weight marker lane, with weights in kDa. (**C**) Immunoblot analysis using anti-PiE1, anti-PilE2 or anti-6His antibodies of sheared filaments that were affinity purified by pull down. The WT strain was included as a control. Sheared filaments were prepared from cultures adjusted to the same OD_600_, affinity purified, eluted in the same final volume and identical volumes were loaded in each lane. MW indicates the molecular weight marker lane, with weights in kDa.

Taken together, these findings show that *S. sanguinis* Tfp are hetero-polymers composed of a 4:3 ratio of PilE1 and PilE2, both harbouring a single PTM, *i.e*. a methylated N-terminus.

### Major pilin processing and/or assembly into filaments in Gram-positive Tfp follow widely conserved principles

Mutagenesis studies of major pilins in several Tfp and/or Tff have identified residues in class III signal peptides key for processing by PilD and/or assembly into filaments (9,29-31). A common rule has emerged concerning two highly conserved residues Gly_-1_ and Glu_5_ (1). Gly_-1_ is crucial for processing by PilD, while Glu_5_ is dispensable for processing but critical for filament assembly (14,15). Since the importance of these residues has not been assessed for Gram-positive Tfp, we tested it in *S. sanguinis*. Because PilE1 and PilE2 have identical N-termini (Fig. 1B and Fig. S2), we focused our efforts on PilE1 and used our gene editing strategy (25) to construct markerless mutant strains expressing PilE1 variants in which the Gly_-1_ and Glu_5_ residues would be mutated. Next, we tested whether these mutants proteins were processed by PilD and assembled into filaments. Pilin processing was assessed by immunoblotting in wholecell protein extracts using the anti-PilE1 antibody (24), *i.e*. processed PilE1 is 14.7 kDa while the unprocessed protein is 16.9 kDa. Assembly into Tfp was assessed by immunoblotting on purified filaments. Originally, we replaced Gly_-1_ by an Ala (PilE1_G-1A_), which had no effect on processing (Fig. 4A) or assembly into filaments (Fig. 4B). A similar finding was reported in *P. aeruginosa*, and was attributed to the small size of Ala since substitutions with bulkier residues abolished pilin processing (9). Accordingly, when Gly_-1_ was replaced by a Ser (PilE1_G-1S_), PilE1 could not be processed (Fig. 4A) or polymerised into filaments, which consisted only of PilE2 (Fig. 4B). The marked decrease in PilE2 in pilus preparation suggests a dominant negative effect of unprocessed PilE1_G-1S_ on PilE2 filament formation. As for the Glu_5_ residue, when it was replaced by an Ala (PilE1_E5A_), PilE1 processing was unaffected (Fig. 4A) but the protein could not be polymerised into filaments, which again consisted only of PilE2 (Fig. 4B). As above, there was a marked decrease in PilE2 in pilus preparation suggesting a dominant negative effect of PilE1_E5A_ on PilE2 filament formation. These findings confirm that the widely conserved principles defining how major pilins are processed and assembled into filaments (1), apply to Gram-positive Tfp as well. This further highlights the suitability of *S. sanguinis* as a Gram-positive model to study Tfp biology.

**Figure 4.**
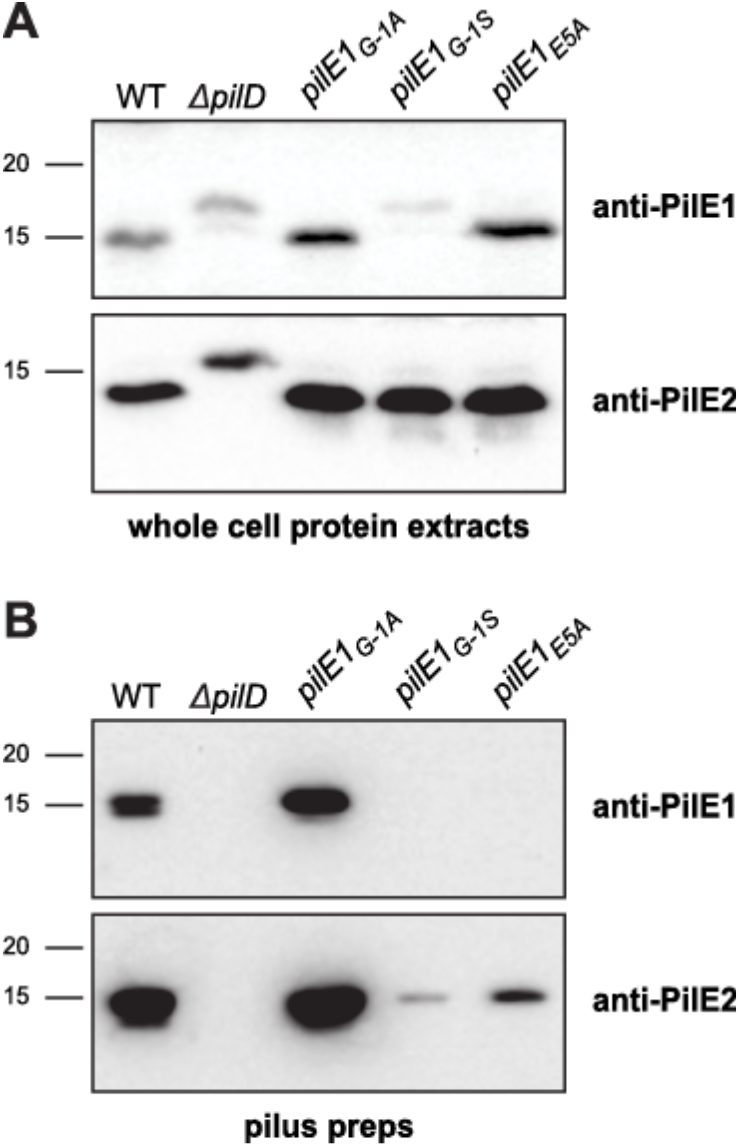
Processing of *S. sanguinis* major pilin PilE1 by PilD and its assembly in filaments follow widely conserved general principles. (**A**) Immunoblot analysis of PilE1 expression and processing by PilD in strains expressing PilE1_G-1A_, PilE1_G-1S_ and PilE1_E5A_ mutants. The WT strain *ΔpilD* mutant have been included as controls. Whole-cell protein extracts were separated by SDS-PAGE and probed using anti-PilE1 antibody or anti-PilE2 antibody as a control. Protein extracts were quantified, equalised and identical volumes were loaded in each lane. Molecular weights are indicated in kDa. (**B**) Immunoblot analysis of PilE1 assembly in filaments in strains expressing PilE1_G-1A_, PilE1_G-1S_ and PilE1_E5A_ mutants. The WT strain and *ΔpilD* mutant have been included as controls. Purified Tfp were separated by SDS-PAGE and probed using anti-PilE1 antibody or anti-PilE2 antibody as a control. Samples were prepared from cultures adjusted to the same OD_600_, and identical volumes were loaded in each lane but to improve detection of faint bands, we loaded 20-fold dilutions of WT and PilE1_G-1A_ samples. Molecular weights are indicated in kDa.

### 3D structure of PilE1 reveals a pilin fold with an uncommon highly flexible C-terminus

Next, to improve our structural understanding of *S. sanguinis* Tfp, we solved the 3D structure of its major pilins. To facilitate purification, we expressed in *Escherichia coli* the soluble portions of PilE1 (112 aa) and PilE2 (105 aa) fused to an N-terminal 6His tag (Fig. S2). This truncated the first 27 residues of the mature proteins, which form the protruding hydrophobic N-terminal α-helix in type IV pilins (1,3). This procedure allowed us to purify well-folded and soluble proteins using a combination of affinity and gel-filtration chromatography. Since PilE2 shares 78% sequence identity with PilE1 (Fig. S2), we decided to focus on the longest PilE1 and determine its structure by NMR. We isotopically labelled 6His-PilE1 with ^13^C and ^15^N for backbone and side-chain NMR resonance assignment (Table S1) and obtained a high-resolution structure in solution. This structure (Fig. 5A) revealed that PilE1 adopts the classical type IV pilin fold (1,3). It exhibits a long N-terminal α-helix packed against a β-meander consisting of three anti-parallel β-strands, flanked by distinctive ‘edges’ (Fig. 5A), which usually differ between pilins (1,3). While the loop connecting α1 and β1 is unexceptional except perhaps for its length (50 aa), the C-terminus of the protein (after β3) is striking. Unlike in other pilins in which this region is stabilised by being ‘stapled’ to the last β-strand either by a disulfide bond, a network of hydrogen bonds, or a calcium-binding site (1,3), the 10 aa-long C-terminus in PilE1 is unstructured and highly flexible. While the structures within the NMR ensemble superpose well up to the last β strand (β3), their C-termini exhibit different conformations and orientations (Fig. 5B). Intriguingly, when compared to structures in the Protein Data Bank (PDB), PilE1 was found to be most similar to pseudopilins, which form short Tff, rather than to *bona fide* Tfp subunits. As can be seen in Fig. 5C, the structures of PilE1 and PulG - the major pseudopilin in *Klebsiella oxytoca* type II secretion system (T2SS) (32) - show extended similarity.

**Figure 5.**
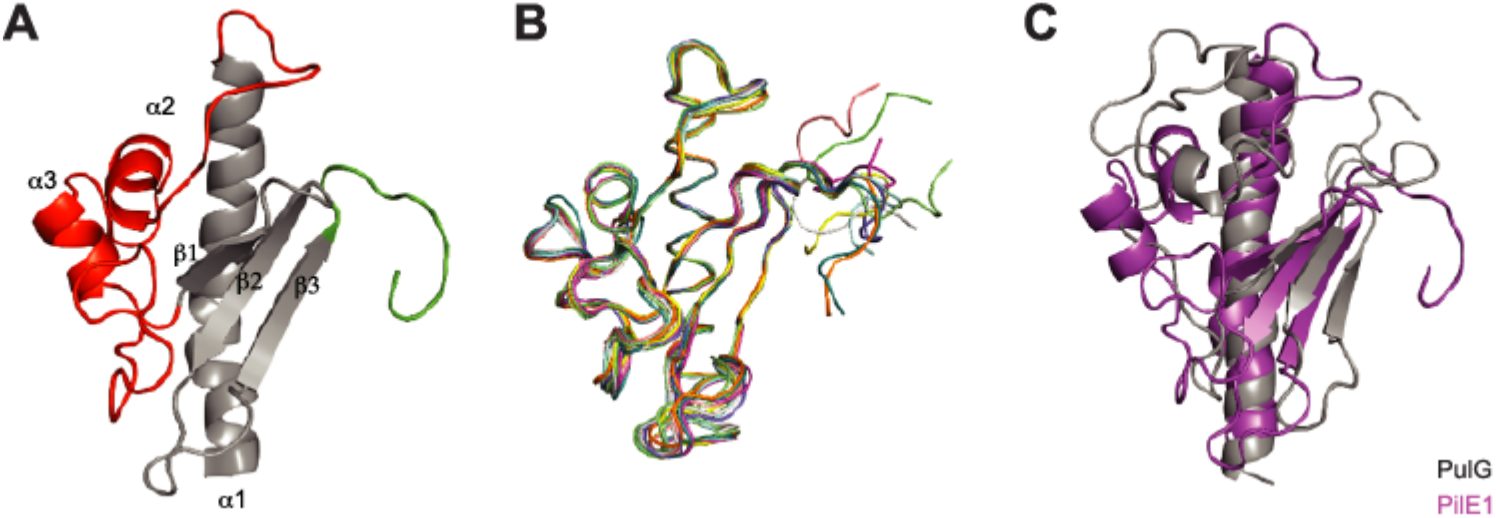
High-resolution 3D structure of PilE1 reveals a canonical type IV pilin fold with an uncommon highly flexible C-terminus, which more closely resembles pseudopilins. (**A**) Cartoon representation of the NMR structure of the globular domain of PilE1. The conserved core in type IV pilins (the N-terminal α-helix and 3-stranded antiparallel β-meander) is depicted in grey. Distinctive/key structural features flanking the β-meander are highlighted in color, the α1β1-loop in red and the unstructured C-terminus in green. (**B**) Ribbon representation of the superposition of the ensemble of 10 most favourable PilE1 structures determined by NMR. (C) Superimposed cartoon representations of the globular domains of PilE1 (magenta) and *K. oxytoca* pseudopilin PulG (grey). The two structures superpose with a rmsd of 5.75 Å over their entire length.

Owing to their high sequence identity (Fig. S2), we used our PilE1 structure as a template to produce a homology model of PilE2 (Fig. 6A). As expected, PilE2 was found to be virtually identical to PilE1 except for its shorter α1β1-loop (Fig. 6B), which is explained by the fact that eight residues at the C-terminus of this loop in PilE1 are absent in PilE2 (Fig. S2). Finally, after producing full-length models of PilE1 and PilE2 using the full-length gonococcal pilin (33,34) as a template, we were able to model packing of these pilins within recently determined structures of Tfpa (14,15) (Fig. 6C). This revealed that PilE1 and PilE2 fit readily into these Tfp, which have a similar morphology to *S. sanguinis* filaments. This finding supports the notion that polymerisation of pilins into filaments in Gram-positive species also occurs via hydrophobic packing of their α1N helix within the filament core.

**Figure 6.**
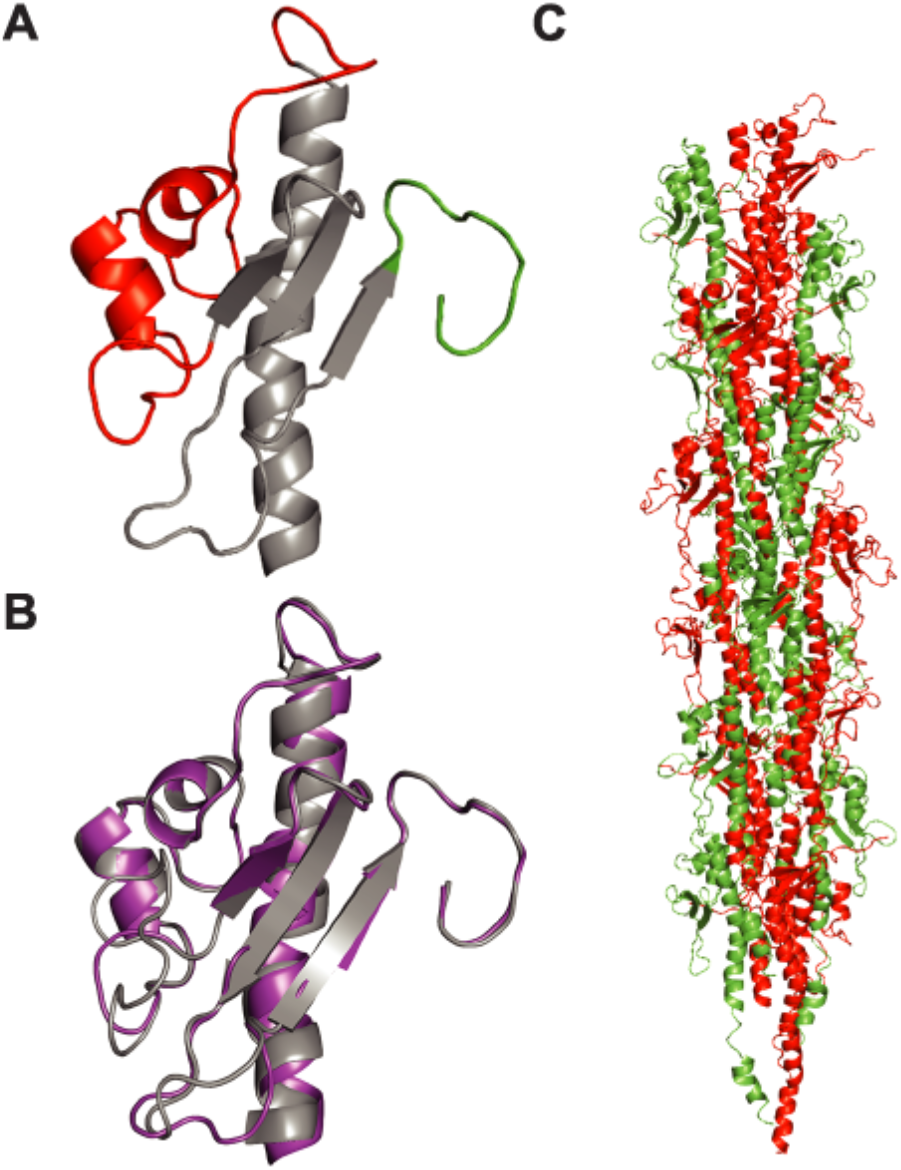
Modelling shows that full-length PilE1/PilE2 fit readily into available Tfp structures. (**A**) Cartoon representation of the homology structural model of PilE2 based on PilE1 structure. Same coulour codes have been used as in Fig. 5A. (**B**) Superposition of globular domains of PilE1 and PilE2 (right) with a rmsd of 0.61 Å. (**C**) Packing of full-length models of PilE1 and PilE2 into the recently determined structure of gonoccoccal Tfp. Half of the gonoccoccal subunits in the structure were replaced by PilE1 and the other half with PilE2.

Together, these findings show that *S. sanguinis* Tfp obey structural principles common to this class of filaments with some intriguing peculiarities. While *S. sanguinis* major pilins adopt a canonical type IV pilin fold, their C-terminus appears to be highly flexible, which is uncommon. Moreover, while *S. sanguinis* major pilins fit readily within available Tfp structures, they are more similar structurally to pseudopilins than to *bona fide* Tfp-forming pilins.

### PilA, PilB and PilC are minor pilin components of S. sanguinis Tfp

In all Tff systems, there are in addition to the major pilins several proteins with class III signal peptides, whose role and localisation are often unclear (3). To start with experimental characterisation of PilA, PilB and PilC in *S. sanguinis*, we purified each protein and generated antisera. Immunoblotting using whole-cell protein extracts confirmed that the three proteins are expressed since they could be detected in WT, but not in the corresponding deletion mutants (Fig. 7A). We then tested whether PilA, PilB and PilC were processed by PilD, which for PilA was uncertain considering its degenerate class III signal peptide (Fig. 1B). Processing by PilD, which removes the leader peptide, is expected to generate mature proteins of 17, 50.5 and 52.8 kDa for PilA, PilB and PilC, respectively, shorter than their 18, 51.5 and 53.9 kDa precursors. Immunoblots confirmed that all three proteins are cleaved by PilD as indicated by the detection of proteins with masses consistent with unprocessed precursors in a *ΔpilD* mutant (Fig. 7A). Next, we took advantage of our ability to prepare highly pure Tfp, to determine whether PilA, PilB and PilC could be detected in pilus preparations by immunoblotting. As can be seen in Fig. 2B, although only PilE1 and PilE2 could be detected by SDS-PAGE/Coomassie analysis, PilA, PilB and PilC are readily detected by immunoblotting in these preparations (Fig. 7B). Co-purification of PilA, PilB and PilC with the filaments was dependent upon their processing by PilD since these proteins were not detected in sheared fractions prepared from a non-piliated *ΔpilD* mutant (Fig. 7B). In conclusion, findings that PilA, PilB and PilC are cleaved by PilD and co-purify with Tfp suggest that these three proteins are minor (low abundance) pilin components of *S. sanguinis* Tfp, likely assembled into filaments in a similar fashion to the major subunits PilE1 and PilE2.

**Figure 7.**
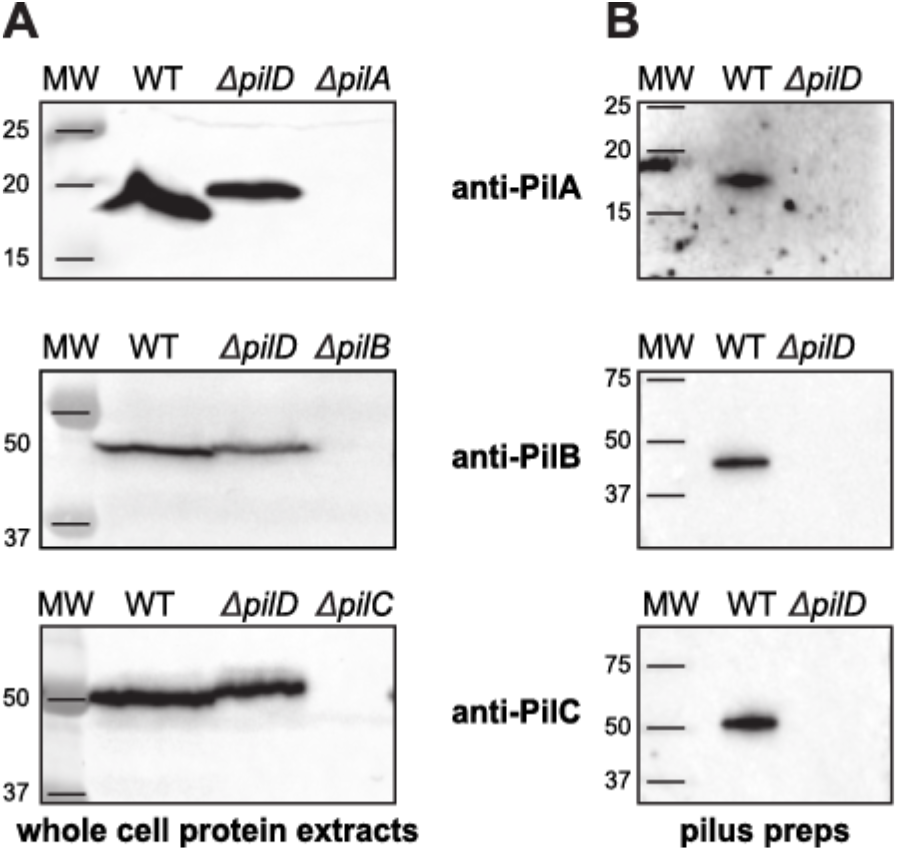
Pilin-like proteins PilA, PilB and PilC are processed by PilD and co-purify with Tfp. (**A**) Immunoblot analysis of PiA, PilB and PilC expression and processing by PilD. Whole-cell protein extracts were probed using specific anti-peptide antibodies, which were generated for this study. Protein extracts were quantified, equalised and equivalent amounts of total proteins were loaded in each lane. Molecular weights are indicated in kDa. (**B**) Immunoblot analysis of pilus preparations using anti-PiA, anti-PilB and anti-PilC antibodies. Samples were prepared from cultures adjusted to the same OD_600_ and identical volumes were loaded in each lane. Molecular weights are indicated in kDa.

### Homo-polymeric filaments composed only of PilE1 or PilE2 are able to promote motility, but at different speeds

As previously reported, while a double *ΔpilE1ΔpilE2* mutant is non-piliated, single *ΔpilE1* and *ΔpilE2* mutants produce WT-like filaments consisting of the remaining pilin (24). We therefore wondered whether these homo-polymeric filaments are functional and, if so, whether the structural differences between the two pilins (see Fig. 6B) would have a functional impact. We compared the ability of the homo-polymeric filaments produced by single *ΔpilE1* and *ΔpilE2* mutants to mediate twitching motility. We first found that these mutants still exhibited spreading zones around bacteria grown on agar plates (Fig. 8A). This confirms that Tfp consisting exclusively of PilE1 or PilE2 are functional. Motility was next assessed quantitatively at a cellular level by tracking under the microscope the movement of small chains of cells (Fig. 8B). As previously reported for the WT (24), both mutants showed ‘train-like’ directional motion mainly parallel to the long axis of bacterial chains. Short duration movies illustrating the movement of *ΔpilE1* and *ΔpilE2* are included as supplementary information (Movies S1 and S2). Measurement of instantaneous velocities revealed that, while the WT moved at 694 ± 4 nm·s^−1^ (mean ± standard error, *n*=22,957) consistent with previous measurements (24), the mutants moved at 462 ± 2 nm·s^−1^ (*n*=37,001) for *ΔpilE1*, and 735 ± 3 nm·s^−1^ (*n*=22,231) for *ΔpilE2* (Fig. 8B). These statistically significant differences suggest that the two pilins are not functionally equivalent with respect to twitching motility.

**Figure 8.**
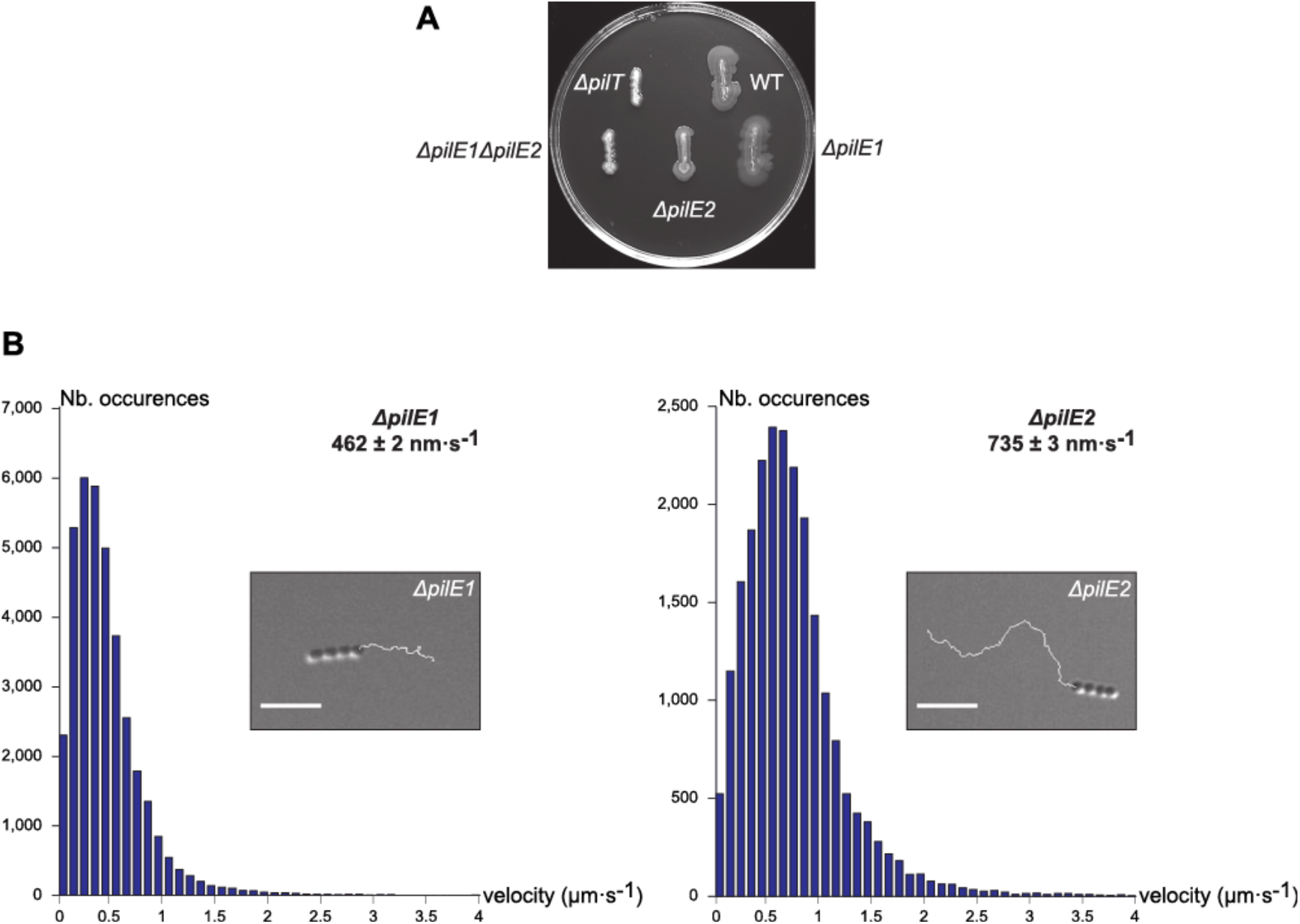
*ΔpilE1* and *ΔpilE2* mutants produce Tfp capable of powering motility, albeit at different speeds. (**A**) Macroscopic motility assay. Spreading zones, or lack thereof, around single and double *ΔpilE1* and *ΔpilE2* mutants. The WT strain and *ΔpilT* mutant have been included as positive and negative controls, respectively. (**B**) Microscopic motility assay. The histograms represents the distribution curve of velocities (in 100 nm·s^−1^ intervals) measured for *ΔpilE1* and *ΔpilE2* mutants. Insets are representative 30 sec trajectories of movement of small chains of cells. Scale bar represents 5 μm. Corresponding movies are available as supplementary information (Movies S1 and S2).

## Discussion

Their ubiquity in prokaryotes (1) makes Tff an important research topic. A better understanding of the molecular mechanisms governing Tff biology might have practical implications for human health and nanotechnology. Perhaps one of the reasons for our limited understanding of Tff biology is that, historically, these filamentous nanomachines have been studied in just a few Gram-negative bacterial species, all belonging to the same phylum (Proteobacteria) (4). The study of Tff in phylogenetically distant species has the potential to move the field forward, which has recently sparked studies in Archaea (2) and in distant phyla of Bacteria (21). One of the most promising new Tfp models that has emerged is the Gram-positive opportunistic pathogen *S. sanguinis* (24,25). A recent systematic genetic analysis of Tfp biology in this species - the first to be realised in a non Proteobacterium - showed that *S. sanguinis* uses a simpler machinery (with fewer components) to assemble canonical Tfpa that generate high tensile forces and power twitching motility (24). In this report, we performed an in depth biochemical and structural analysis of *S. sanguinis* filaments, which led to the notable findings discussed below.

The first important achievement in this study is the etablishment of one of the most complete biochemical pictures of a Tfp. Criticallly, this confirms the above observed trend (24) that *S. sanguinis* Tfpa are simpler filaments, with fewer components. Indeed, in Gram-negative Tfpa models, in addition to the major pilin there are 7-8 proteins possessing class III signal peptides (4). For example, in *Neisseria meningitidis*, there are four conserved pilin-like proteins required for piliation (PilH, PilI, PilJ and PilK) whose role (priming filament assembly) (35) and localisation (at the tip of the pili or distributed throughout the filaments) (36,37) remain uncertain, and three species-specific minor pilins (ComP, PilV and PilX) that are dispensable for piliation but modulate Tfp-associated functions (38-40). In contrast, in *S. sanguinis*, besides PilE1 and PilE2 there are only three Pil proteins possessing class III signal peptides (PilA, PilB and PilC). As shown here, these proteins are efficiently recognised and post-translationally modified by PilD, which processes their N-terminal leader peptides and (most likely) methylates the first residue of the resulting mature proteins, as demonstrated for PilE1 and PilE2. Processing follows widely conserved principles (1). No other PTM, which frequently decorate major pilins in Gram-negative Tfp are found on *S. sanguinis* PilE1 and PilE2. Importantly, the use of highly pure pilus preparations clearly showed that these five proteins are Tfp subunits. PilE1 and PilE2 are the two major pilins, while PilA, PilB and PilC are three minor pilins. Although the arrangement of the major pilins in the filaments (geometric or stochastic) remains to be determined, this study strongly suggests that *S. sanguinis* Tfp are hetero-polymeric structures containing comparable amounts of PilE1 and PilE2, a property not previously reported for Tff. However, the reason for this peculiarity remains unclear since the homo-polymeric Tfp assembled by *ΔpilE1* and *ΔpilE2* mutants power efficient twitching motility. A possible explanation might be that PilE1 and PilE2 are important for optimal stability of *S. sanguinis* Tfp as *ΔpilE1* and *ΔpilE2* mutants produce less filaments than WT. On the other hand, what could be the role of the minor subunits of *S. sanguinis* Tfp (PilA, PilB and PilC)? Although they are required for piliation (24), they are unlikely to prime filament assembly because they are unrelated to the four conserved pilin-like proteins carrying out this process in Gram-negative Tfp (35). Rather, PilA, PilB and PilC are likely to contribute to filament stability and to modulate Tfp-associated functions. This notion is supported by the unusual presence of large C-terminal domains in PilB and PilC. Interestingly, both the von Willebrand factor type A motif (found in PilB) and the concanavalin A-like lectin/glucanase structural domain (found in PilC) are often involved in binding mammalian protein or carbohydrate ligands, suggesting that PilB and PilC are involved in host adhesion in *S. sanguinis*, a property frequently associated with Tfp in other species (4).

The structural information generated on *S. sanguinis* Tfp is the second notable achievement in this study. Our new pilus purification strategy shows that the morphological features of *S. sanguinis* filaments are canonical of Tfpa (4). It is possible that the thick and irregular filaments purified previously (24) were damaged during ultra-centrifugation by the presence of cells/cellular debris. Our high-resolution NMR structure of the globular domain of PilE1 confirms that Gram-positive Tfp major subunits adopt the classical type IV pilin fold (1,3). The 3D structure of PilE2 is likely to be virtually identical, owing to its ~80% sequence identity to PilE1. Full-length PilE1/PilE2 are therefore expected to adopt the canonical ‘lollipop’ structure (33,34), since the missing hydrophobic α1N portion can reliably be modelled as a protruding α-helix. Their α1 helix is likely to adopt a gentle S-shaped curve like in the gonococcal major pilin (33,34), because the helix-breaking Pro22 is conserved. Importantly, PilE1/PilE2 fit well in recent cryo-electron microscopy (cryo-EM) reconstructions of several Gram-negative Tfpa (14,15), which despite different helical parameters display similar packing of the α1N helix within the core of the filaments. Owing to the conservation of the helix-breaking Pro22, it is likely that the α1N helix will be partially melted between Ala14 and Pro_22_ as the pilins in the above reconstructions (14,15), which is thought to provide flexibility and elasticity to the filaments (41). In addition, this would allow the formation of a salt bridge between Glu_5_ and the methylated Phe_1_ (both conserved in PilE1 and PilE2) of the neighbouring pilin. Minor subunits PilB and PilC, which have canonical class III signal peptides similar to PilE1/PilE2 are likely to assemble within filaments in a similar fashion. It is unclear, however, whether the α1N helix would be partially melted since the helix-breaking Pro_22_ is absent in these proteins. Moreover, it remains to be seen how the bulky C-terminal domains in PilB and PilC, which appear to have been ‘grafted’ by evolution onto a pilin moiety, would be exposed on the surface of the filaments. As for PilA, its unique class III signal peptide, makes it difficult to predict how its α1N helix will be packed in the filament core, especially because of the unusual Pro_4_. Critically, our high-resolution structure of PilE1 also challenges two common assumptions in the field (1,3). First, in contrast to all previously available pilin structures, the C-terminus in PilE1 is unstructured and highly flexible. This explains why it is a permissive insertion site and why it even can be deleted and replaced by a 6His tag (25), without interfering with the ability of PilE1/PilE2 to be polymerised into filaments. Therefore, the common rule that the C-terminus of major pilins must be stabilised to preserve pilin integrity and its ability to be polymerised (3), is not always true. It cannot be excluded at this point that the major pilins of *S. sanguinis* are an exception as the major pilin of another Tfp-expressing Gram-positive, *Clostridium difficile*, apparently needs to stabilise its C terminus (42). Second, although a Tfp-forming major pilin, PilE1 is most similar to pseudopilins that form short filaments in alternative Tff, such as PulG from *K. oxytoca* T2SS (32). This indicates that a major pilin 3D structure cannot be used to predict whether the subunit will form pseudopili or *bona fide* Tfp, which perhaps blurs the lines between different Tff and suggests an even closer evolutionary relationship between these filamentous nanomachines.

In conclusion, by providing an unprecedented global view of a Gram-positive Tfp, this study further cements *S. sanguinis* as a model species which is fast closing the gap with historic Gram-negative Tfp models. Together with our recent reports (24,25) and *S. sanguinis* exquisite genetic tractability, these findings pave the way for future investigations, which will undoubtedly contribute to improve our understanding of a fascinating filamentous nano-machine almost universal in prokaryotes.

### Experimental procedures

#### Strains and growth conditions

Strains and plasmids that were used in this study are listed in Table S2. For cloning, we used *E. coli* DH5α. *E. coli* BL21(DE3) was used for protein purification. *E. coli* strains were grown in liquid or solid Lysogenic Broth (LB) (Difco) containing, when required, 100 μg/ml spectinomycin or 50 μg/ml kanamycin (both from Sigma). The WT *S. sanguinis* 2908 strain and deletion mutants were described previously (24,25). *S. sanguinis* strains were grown on plates containing Todd Hewitt (TH) broth (Difco) and 1% agar (Difco), incubated at 37°C in anaerobic jars (Oxoid) under anaerobic conditions generated using Anaerogen sachets (Oxoid). Liquid cultures were grown statically under aerobic conditions in THT, *i.e*. TH broth containing 0.05% tween 80 (Merck) to limit bacterial clumping. When required, 500 μg/ml kanamycin (Km) (Sigma) was used for selection. For counterselection, we used 15 mM *p*-Cl-Phe (Sigma) (25).

Chemically competent *E. coli* cells were prepared as described (43). DNA manipulations were done using standard molecular biology techniques (44). All PCR were done using high-fidelity DNA polymerases from Agilent (see Table S3 for a list of primers used in this study). *S. sanguinis* genomic DNA was prepared from overnight (O/N) liquid cultures using the kit XIT Genomic DnA from Gram-Positive Bacteria (G-Biosciences). Strain 2908, which is naturally competent, was transformed as described elsewhere (24,25).

Unmarked *S. sanguinis* mutants in *pilE1* and *pilE2 in situ* used in this study were constructed using a recently described two-step, cloning-independent, gene editing strategy (25). In brief, in the first step, the target gene was cleanly replaced in the WT, by allelic exchange, with a promoterless *pheS*aphA-3* double cassette, which confers sensitivity to *p*-Cl-Phe and resistance to kanamycin. To do this, a splicing PCR (sPCR) product fusing the upstream (amplified with primers F1 and R1) and downstream (amplified with F2 and R2) regions flanking the target gene to *pheS*aphA-3* (amplified with *pheS*-F and *aph*-R) was directly transformed into the WT, and allelic exchange mutants were selected on Km-containing plates. Allelic exchange was confirmed for a couple of transformants by PCR. In the second step, the *pheS*aphA-3* double cassette was cleanly replaced in this primary mutant, by allelic exchange, with an unmarked mutant allele of the target gene (see below). To do this, an sPCR product fusing the mutant allele to its upstream and downstream flanking regions was directly transformed into the primary mutant and allelic exchange mutants were selected on *p*-Cl-Phe-containing plates. Markerless allelic exchange mutants, which are Km^S^, were identified by re-streaking *p*-Cl-Phe^R^ colonies on TH plates with and without Km. The *pilE1* and *pilE2* mutant alleles encoding proteins with a C-terminal 6His tag were engineered by sPCR. We made two constructs for each gene: a LONG construct in which we fused the 6His tag to the C-terminus of the full-length protein (sPCR with F1/R3 and F3/R2), and a SHORT construct in which we replaced the last seven aa by the tag (sPCR with F1/R4 and F4/R2). To construct the missense mutants in *pilE1*, we used as a template a pCR8/GW/TOPO plasmid in which the WT gene (amplified with F and R) was cloned (Table S2) and the Quickchange site-directed mutagenesis kit (Agilent) (with complementary primers #1 and #2). Then, the sPCR product for transformation in the primary mutant was produced by fusing the mutant allele (amplified with F and R) to flanking regions upstream (amplified with F1 and R5) and downstream (amplified with F5 and R2).

#### SDS-PAGE, antisera and immunoblotting

*S. sanguinis* whole-cell protein extracts were prepared using a FastPrep-24 homogeniser (MP Biomedicals) and quantified as described elsewhere (24). Separation of the proteins by SDS-PAGE, subsequent blotting to Amersham Hybond ECL membrane (GE Healthcare) and blocking were carried out using standard molecular biology techniques (44). To detect PilE1 and PilE2, we used previously described primary rabbit anti-peptide antibodies (24). Antisera against PilA, PilB and PilC were produced for this study by Eurogentec by immunising rabbits with purified recombinant proteins (see below). These proteins were then used to affinity-purify the antibodies. Primary antibodies were used at between 1/2,000 and 1/5,000 dilutions, while the secondary antibody, an ECL HRP-linked anti-rabbit antibody (GE Healthcare), was used at 1/10,000 dilution. Amersham ECL Prime (GE Healthcare) was used to reveal the blots. To detect His-tagged proteins, we used a HRP-linked anti-6His antibody (Sigma) at 1/10,000 dilution.

#### Tfp purification and visualisation

*S. sanguinis* 2908 Tfp were purified as described elsewhere with minor modifications (24). Liquid cultures (10 ml), grown O/N in THT, were used the next day to re-inoculate 90 ml of THT and grown statically until the OD_600_ reached 1-1.5, at which point OD were normalised, if needed. Bacteria were pelleted at 4°C by centrifugation for 10 min at 6,000 *g* and pellets were re-suspended in 2 ml pilus buffer (20 mM Tris pH 7.5, 50 mM NaCl). This suspension was vortexed for 2 min at full speed to shear Tfp. Bacteria were then pelleted as above, and supernatant containing the pili was transferred to a new tube. This centrifugation step was repeated, before the supernatant was passed through a 0.22 μm pore size syringe filter (Millipore) to remove residual cells and cellular debris. Pili were then pelleted by ultra-centrifugation as described (24), resuspended in pilus buffer, separated by SDS-PAGE and gels were stained using Bio-Safe Coomassie stain (Bio-Rad). Purified filaments were visualised by TEM after negative staining as described elsewhere (24).

Pull down purification of 6 His tagged filaments was done as follows. Fifty μl of Dynabeads His-Tag isolation and pull down beads (Life Technologies) were aliquoted in an Eppendorf tube, and pelleted by placing the tube on a magnet for 2 min. After decanting the buffer, beads were washed three times using 500 μl of pilus buffer, mixed with 900 μl of sheared filaments (prepared as for the purification above) and incubated for 10 min on a roller at room temperature. Then, the tube was placed on the magnetic holder for 2 min and the supernatant discarded. Beads were subsequently washed three times with 400 μl of pilus buffer, and finally incubated for 10 min with 100 μl of elution buffer (pilus buffer with 500 mM imidazole) on a roller at room temperature. Then, the tube was placed on the magnetic holder for 2 min, the supernatant collected and analysed by SDS-PAGE/Coomassie and/or immunoblotting.

#### Proteomics and mass spectrometric analysis of purified Tfp

For the bottom-up MS analysis of purifed Tfp, we carefully excised PilE1 and PilE2 protein bands from Coomassie-stained gels and generated enzymatically derived peptides employing four separate enzymes. In brief, as previously described (45), gel pieces were destained and digested O/N with 1 μg trypsin or Lys-C in (50 mM Na_2_HCO_3_, pH 7.8) at 37°C, 1 μg chymotrypsin in (100 mM Tris(hydroxymethyl)aminomethane, 10 mM CaCl_2_·2H_2_O, pH 7.8) at 25°C, or 0.3 μg AspN in (50 mM Tris(hydroxymethyl)aminomethane, 2.5 mM ZnSO_4_·7H_2_O, pH 8.0) at 25°C. Generated peptides, which were extracted as previously described (45), were vacuum-concentrated, dissolved in loading buffer (2% acetonitrile, 1% trifluoroacetic acid) and desalted using ZIP-TIP tips as instructed by the manufacturer (Millipore). The peptides were eluted with (80% acetonitrile, 1% trifluoroacetic acid) and vacuum-concentrated. Dried peptide samples were dissolved in 10 μl (2% acetonitrile, 1% formic acid) before analysis by reverse phase LC-MS/MS. Redissolved samples (2-5 μl) were injected into a Dionex Ultimate 3000 nano-UHPLC system (Sunnyvale) coupled online to a QExactive mass spectrometer (Thermo Fisher Scientific) equipped with a nano-electrospray ion source. LC separation was achieved with an Acclaim PepMap 100 column (C18, 3 μm beads, 100 Å, 75 μm inner diameter, 50 cm) and a LC-packing trap column (C18, 0.3 mm inner diameter, 300 Å). The flow rate (15 μl/min) was provided by the capillary pump. A flow rate of 300 nl/min was employed by the nano pump, establishing a solvent gradient of solvent B from 3 to 5% in 5 min and from 5 to 55% in 60 min. Solvent A was (0.1% formic acid, 2% acetonitrile), while solvent B was (0.1% formic acid, 90% acetonitrile). The mass spectrometer was operated in data-dependent mode to automatically switch between MS and MS/MS acquisition. Survey full scan MS spectra (from *m/z* 200 to 2,000) were acquired with the resolution R = 70,000 at *m/z* 200, with an automated gain control (AGC) target of 10^6^, and ion accumulation time set at 100 msec. The seven most intense ions, depending on signal intensity (intensity threshold 5.6×10^3^) were considered for fragmentation using higher-energy collisional induced dissociation (HCD) at R = 17,500 and normalised collision energy (NCE)=30. Maximum ion accumulation time for MS/MS spectra was set at 180 msec. Dynamic exclusion of selected ions for MS/MS were set at 30 sec. The isolation window (without offset) was set at *m/z* 2. The lock mass option was enabled in MS mode for internal recalibration during the analysis.

To perform a complete top-down MS analysis of the PilE1 and PilE2 proteoforms present in purified *S. sanguinis* Tfp, three separate methods were employed. For the first method, sample preparation and top-down ESI-MS on a LTQ Orbitrap were performed as previously described (46), and intact protein mass spectra were acquired with a resolution of 100,000 at *m/z* 400. For the second method, after initial preliminary testing of a gradient of solvent B from 3 to 55% in 10 min, and from 55 to 85% in 12-35 min, the data was acquired on an LTQ Orbitrap operated in positive ionisation mode in the data-dependent mode, to automatically switch between MS and MS/MS acquisition. Survey full scan MS spectra (from *m/z* 200 to 2,000) were acquired with a resolution R = 100,000 at *m/z* 400, with AGC target of 10^6^ and ion accumulation time set at 100 msec. The two most intense ions, depending on signal intensity (intensity threshold 5.6×10^3^) were considered for fragmentation using HCD at R = 17,500 and NCE=30. The lock mass option was enabled in MS mode for internal recalibration during the analysis. For the third method, a direct injection nano-ESI top-down procedure was employed. Briefly, the Dionex Ultimate 3000 nano-UHPLC system coupled to the LTQ Orbitrap was reconfigured so that after sample loading onto the loop, valve switching allowed the nanopump to inject directly the sample from the loop into the LTQ Orbitrap MS using an isocratic gradient of (10% methanol, 10% formic acid). Data acquisition was done manually from the LTQ Tune Plus (V2.5.5 sp2) using R=100,000 and NCE=400.

Data processing and analysis was done as follows. Bottom-up MS data were analysed using MaxQuant (v 1.5.2) and the Andromeda search engine against an in-house generated *S. sanguinis* whole proteome database, and a database containing common contaminants. Trypsin, chymotrypsin, Lys-C, AspN and no enzyme (no restriction) were selected as enzymes, allowing two missed cleavage sites. We applied a tolerance of 10 ppm for the precursor ion in the first search, 5 ppm in the second, and 0.05 Da for the MS/MS fragments. In addition to methionine oxidation, protein N-terminal methylation was allowed, in a separate search, as a variable modification. The minimum peptide length was set at 4 aa, and the maximum peptide mass at 5.5 kDa. False discovery rate was set at <0.01. Deconvolution of the PilE1 and PilE2 mass envelopes from top-down analysis was done as previously described (47). Deconvoluted protein masses are reported as monopronated [M+H+]. Theoretical masses of PilE1 and PilE2 were determined from available sequences.

#### Twitching motility assays

Twitching motility was assessed macroscopically on agar plates as described elsewhere (24). Briefly, bacteria were re-streaked as straight lines on freshly poured TH plates containing 1% Eiken agar (Eiken Chemicals), which were incubated up to several days under anaerobic conditions in a jar, in the presence of water to ensure high humidity. Motility was analysed microscopically as described elsewhere (24). In brief, bacteria resuspended in THT were added into an open experimental chamber with a glass bottom and grown for 2 h at 37°C in presence of 5% CO_2_. The chamber was then transferred to an upright Ti Eclipse microscope (Nikon) with an environment cabinet maintaining the same growth conditions, and movies of the motion of small bacterial chains were obtained and analysed in ImageJ, as described (24). Cell speed was measured from collected trajectories using Matlab.

#### Protein purification

To produce pure PilA, PilB and PilC proteins for generating antibodies, we cloned the corresponding genes in pET-28b (Novagen) (Table S2). The forward primer was designed to fuse a non-cleavable N-terminal 6His tag to the soluble portion of these proteins, *i.e*. excluding the leader peptide and the predicted hydrophobic α1N helix. For *pilA*, we amplified the gene from 2908 genome, while for *pilB* and *pilC*, we used synthetic genes (GeneArt), codon-optimised for expression in *E. coli*. Recombinant proteins were purified using a combination of affinity and gel-filtration chromatographies as follows. An O/N liquid culture, in selective LB, from a single colony of *E. coli* BL21 (DE3) transformed with the above expression plasmids, was back-diluted (1/500) the next day in 1 l of the same medium and grown to an OD_600_ of 0.4-0.6 on an orbital shaker. The temperature was then set to 16°C, the culture allowed to cool for 30 min, before protein expression was induced O/N by adding 0.1 mM IPTG (Merck Chemicals). The next day, cells were harvested by centrifugation at 8,000 *g* for 20 min and subjected to one freeze/thaw cycle in binding buffer A [50 mM HEPES pH 7.4, 200 mM NaCl, 10 mM imidazole, 1 x SIGMAFAST EDTA-free protease inhibitor cocktail (Sigma)]. Cells were disrupted by repeated cycles of sonication, *i.e*. pulses of 5 sec on and 5 sec off during 3-5 min, until the cell suspension was visibly less viscous. The cell lysate was then centrifuged for 30 min at 17,000 *g* to remove cell debris. The clarified lysate was then mixed with 2 ml of Ni-NTA agarose resin (Qiagen), prewashed in binding buffer A, and incubated for 2 h at 4°C with gentle agitation. This chromatography mixture was then filtered through a Poly-Prep gravity-flow column (BioRad) and washed several times with binding buffer A, before the protein was eluted with elution buffer A (50 mM HEPES pH 7.4, 200 mM NaCl, 500 mM imidazole, 1 x SIGMAFAST EDTA-free protease inhibitor cocktail). The affinity-purified proteins were further purified, and simultaneously buffer-exchanged into (50 mM HEPES pH 7.4, 200 mM NaCl), by gel-filtration chromatography on an Akta Purifier using a Superdex 75 10/300 GL column (GE Healthcare).

For structural characterisation of PilE1, we cloned, as above, the portion of *pilE1* encoding the soluble portion of this protein in pET-28b in order to produce a protein with a non-cleavable N-terminal 6His tag (Table S2). An O/N pre-culture in LB was back-diluted 1/50 into 10 ml selective M9 minimal medium, supplemented with a mixture of vitamins and trace elements. This was grown to saturation O/N at 30°C in an orbital shaker, then back-diluted 1/500 into 1 l of the same medium containing ^13^C D-glucose and ^15^N NH_4_Cl for isotopic labelling. Cells were grown in an orbital shaker at 30°C until the OD_600_ reached 0.8, then 0.4 mM IPTG was added to induce protein production O/N at 30°C. As above, cells were then harvested and disrupted in binding buffer B (50 mM Tris-HCl pH 8.5, 200 mM NaCl, 10 mM imidazole, 1 x SIGMAFAST EDTA-free protease inhibitor cocktail). The protein was first purified by affinity chromatography and eluted in (50 mM Tris-HCl pH 8.5, 200 mM NaCl, 200 mM imidazole, 1 x SIGMAFAST EDTA-free protease inhibitor cocktail). It was then further purified and buffer-exchanged into (50 mM Na_2_HPO_4_/NaH_2_PO_4_ pH 6, 200 mM NaCl) by gel-filtration chromatography using a Superdex 75 10/300 GL column.

#### NMR structure determination of PilE1

Structure determination of PilE1 was done by NMR, essentially as described (48). In brief, isotopically labelled purified 6His-PilE1 was concentrated to ~750 μM in NMR buffer (50 mM Na_2_HPO_4_/NaH_2_PO_4_ pH 6, 50 mM NaCl, 10% D2O). A full set of triple resonance NMR spectra was recorded on a Bruker Avance III 800 MHz spectrometer equipped with triple resonance cryoprobes at 295 K, and processed with NMRPipe (49). Backbone assignments were completed using a combination of HBHA, HNCACB, HNCO, HN(CA)CO, and CBCA(CO)NH experiments using NMRView (One Moon Scientific) (50). Side-chain resonance assignments were obtained from a combination of CC(CO)NH, HC(C)H-TOCSY and (H)CCH-TOCSY experiments using an inhouse software developed within NMRView (51). Distance restraints were obtained from 3D ^1^H^1^H^15^N-NOESY and ^1^H^1^H^13^C-NOESY spectra and used for structure calculations in ARIA 2.3 (52), along with dihedral angle restraints obtained from chemical shift values calculated using the TALOS+ server (53). For each round of six calculations, 100 structures were calculated over eight iterations. In the final iteration, the 10 lowest energy structures were submitted to a water refinement stage to form the final structural ensemble.

#### Bioinformatics

All the sequences were from the genome of *S. anguinis* 2908 (24). Protein alignments were done using the Clustal Omega server at EMBL-EBI, with default parameters. Pretty-printing and shading of alignment files was done using BOXSHADE server at ExPASy. Prediction of functional domains was done by scanning the protein sequence either against InterPro protein signatures (26), or against the SUPERFAMILY database of structural protein domains (28). Both analyses were done using default parameters. MODELLER was used for modelling protein 3D structures (54). In brief, a homology model for full-length PilE1 was produced using MODELLER in multiple template mode, with the N-terminal 25 residues of PilE from the gonococcal Tfp (15) and a representative of the PilE1 NMR ensemble determined in this study. This PilE1 model was then used as a template to produce a PilE2 model using MODELLER in single template mode. These two molecules were then superimposed onto a single chain each of the gonococcal filament cryo-EM reconstruction (15), using COOT (55) using the SSM superpose function. Sections of the polypeptide were then refined into the electron densities using the real space refine function in COOT to adjust fitting of PilE1/PilE2 monomers into the helical filament. Multiple copies of these models were then used to produce the final representation. Structural homologs of PilE1 were identified by scanning the Protein Data Bank (PDB) using the Dali server (56).

## Acknowledgments

We are grateful to Sophie Helaine (Imperial College London), Chiara Recchi (Imperial College London) and Romé Voulhoux (CNRS Marseille) for critical reading of this manuscript. We would like to express our gratitude to Christian Köhler (University of Oslo) for his valuable input and assistance in re-configuring the LC.

## Accession numbers

The NMR solution structure of PilE1 has been deposited in the Worldwide Protein Data Bank under ID code 6I2O and in the Biological Magnetic Resonance Data Bank under ID code 34325.

## Footnotes

This work was supported by grants from the Medical Research Council (MRC) to VP (MR/L008408/1 and MR/P022197/1). IG was a recipient of a PhD studentship from the MRC Centre of Molecular Bacteriology and Infection.

## The abbreviations used are

Tfp: , Type IV pili;
Tff: , type IV filaments;
NMR: , nuclear magnetic resonance;
PTM: , post-translational modifications;
MS: , mass spectrometry;
WT: , wild-type;
PDB: , Protein Data Bank;
T2SS: , type II secretion system;
TH: , Todd Hewitt;
LB: , Lysogenic Broth;
TEM: , transmission electron microscopy;
cryo-EM: , cryo-electron microscopy;
O/N: , overnight;
sPCR: , splicing PCR;
AGC: , automated gain control;
HCD: , higher-energy collisional induced dissociation;
NCE: , normalised collision energy

