## Supplemental information for "Global biochemical and structural analysis of the type IV pilus from the Gram-positive bacterium *Streptococcus sanguinis*"

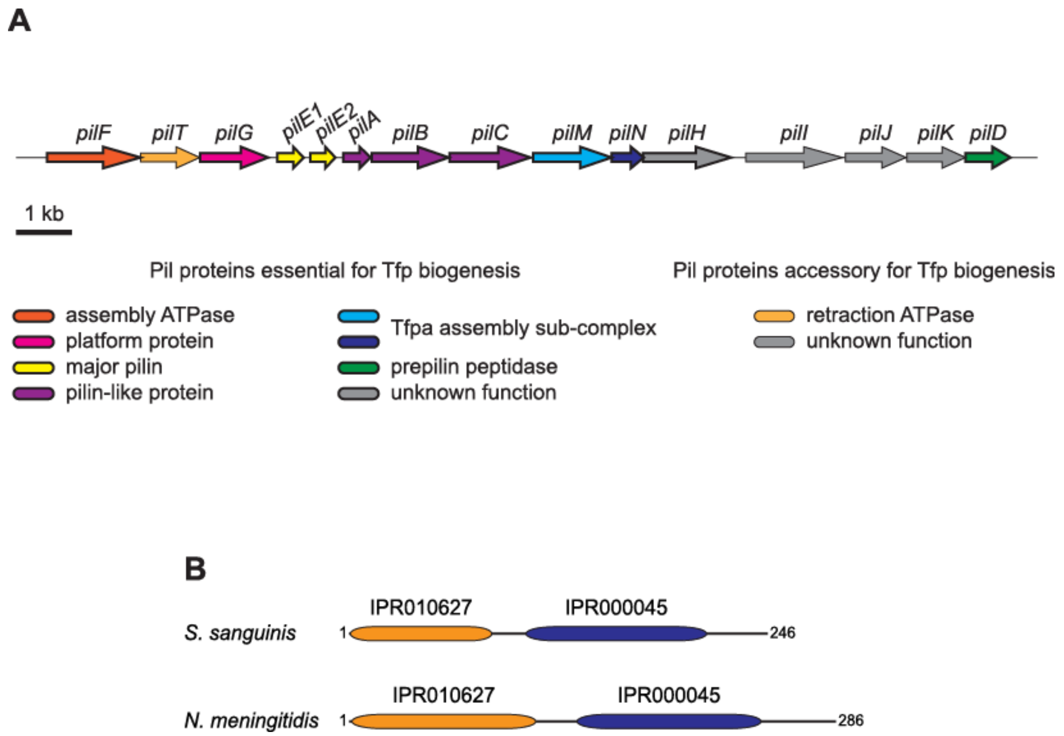


**Figure S1.** Bioinformatic analysis of the genes in the *pil* locus of *S. sanguinis* 2908 and of its prepilin peptidase PilD. (**A**) Gene organisation in the *pil* locus in *S. sanguinis* 2908. All the genes are drawn to scale and the scale bar represents 1 kb. Genes essential for Tfp biogenesis are boxed by a thick line. (**B**) Protein architecture of the prepilin peptidases in *S. sanguinis* and *N. meningitidis*. The N-terminal IPR010627 motif (orange rounded rectangle) catalyses N-methylation, while the C-terminal IPR000045 motif (blue rounded rectangle) catalyses proteolytic processing of pilins. Proteins have been drawn to scale and the subscript numbers indicate protein length.

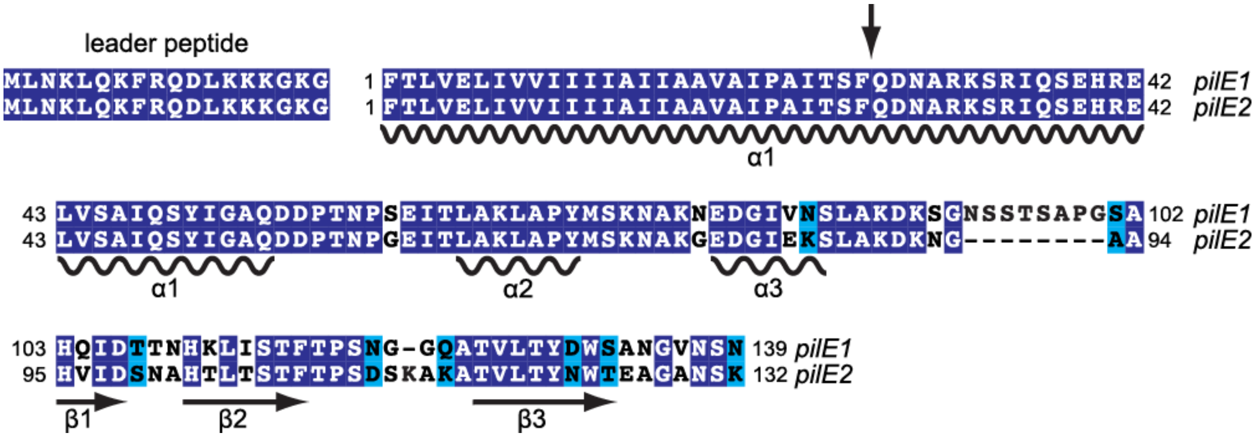


**Figure S2.** Sequence alignment of the major pilins PilE1 and PilE2 in *S. sanguinis* 2908. Residues are shaded in dark blue (identical), light blue (conserved) or unshaded (different). Structural features - α-helices and β-strands - are indicated below the sequences. The vertical arrow indicates the N-terminal portion of PilE1 and PilE2 that was truncated to facilitate protein purification.

**Movies S1.** Cellular motility of a *ΔpilE1* mutant. A small chain of cells attached to a coverslip was imaged for 30 sec. The scale bar represents 5 μm.

**Movies S2.** Cellular motility of a *ΔpilE2* mutant. A small chain of cells attached to a coverslip was imaged for 30 sec. The scale bar represents 5 µm.

**Table S1.** NMR structural statistics.

| | |
|---|---|
| **6His-PilE1** | |
| Number of distance restraints | 1,750 |
| intra-residual | 712 |
| sequential | 416 |
| medium range | 287 |
| long range | 335 |
| NOE violations >0.5 Å (%) | 0.85 |
| Dihedral violations >5º (%) | 0 |
| Ramachandran favoured (%) | 84.4 |
| Ramachandran allowed (%) | 13.5 |
| Ramachandran generously allowed (%) | 0.7 |
| Ramachandran disallowed (%) | 1.3 |

**Table S2.** Strains and plasmids used in this study.

| Name | Details | Source |
|---|---|---|
| ***E. coli* strains** | | |
| DH5α | used for cloning | |
| BL21(DE3) | used for protein expression and purification | |
| ***S. sanguinis* strains** | | |
| 2908 | sequenced WT isolate | (1) |
| *ΔpilA* | *ΔpilA::aphA-3* deletion mutant | (1) |
| *ΔpilB* | *ΔpilB::aphA-3* deletion mutant | (1) |
| *ΔpilC* | *ΔpilC::aphA-3* deletion mutant | (1) |
| *ΔpilD* | *ΔpilD::aphA-3* deletion mutant | (1) |
| *ΔpilE1* | *ΔpilE1::aphA-3* deletion mutant | (1) |
| *ΔpilE1* primary mutant | *ΔpilE1::pheS*aphA-3* deletion mutant | (2) |
| $pilE1_{G\text{-}1A}$ | *pilE1* point mutant expressing $PilE1_{G\text{-}1A}$ | this study |
| $pilE1_{G\text{-}1S}$ | *pilE1* point mutant expressing $PilE1_{G\text{-}1S}$ | this study |
| $pilE1_{E5A}$ | *pilE1* point mutant expressing $PilE1_{E5A}$ | this study |
| $pilE1_{6His\text{-}short}$ | *pilE1* point mutant expressing 6His-tagged $PilE1_{short}$ | this study |
| $pilE1_{6His\text{-}long}$ | *pilE1* point mutant expressing 6His-tagged $PilE1_{long}$ | (2) |
| *ΔpilE2* | *ΔpilE2::aphA-3* deletion mutant | (1) |
| *ΔpilE2* primary mutant | *ΔpilE2::pheS*aphA-3* deletion mutant | this study |
| $pilE2_{6His\text{-}short}$ | *pilE2* point mutant expressing 6His-tagged $PilE2_{short}$ | this study |
| $pilE2_{6His\text{-}long}$ | *pilE2* point mutant expressing 6His-tagged $PilE2_{long}$ | this study |
| *ΔpilE1ΔpilE2* | *ΔpilE1ΔpilE2::aphA-3* double deletion mutant | (1) |
| *ΔpilT* | *ΔpilT::aphA-3* deletion mutant | (1) |
| **Plasmids** | | |
| pCR8/GW/TOPO | TA cloning vector | Invitrogen |
| TOPO-*pheS*aphA-3* | *pheS*aphA-3* double cassette in pCR8/GW/TOPO | (2) |
| TOPO-*pilE1* | full-length *pilE1* in pCR8/GW/TOPO | this study |
| TOPO-$pilE1_{G\text{-}1A}$ | $pilE1_{G\text{-}1A}$ in pCR8/GW/TOPO | this study |
| TOPO-$pilE1_{G\text{-}1S}$ | $pilE1_{G\text{-}1S}$ in pCR8/GW/TOPO | this study |
| TOPO-$pilE1_{E5A}$ | $pilE1_{E5A}$ in pCR8/GW/TOPO | this study |
| pMK-*pilB* | codon-optimised *pilB* in pMK | GeneArt |
| pMK-RQ-*pilC* | codon-optimised *pilC* in pMK-RQ | GeneArt |
| pET-28b | T7-based expression vector | Novagen |
| pET28-*pilA* | pET-28b derivative for expressing 6His-$PilA_{33\text{-}164}$ | this study |
| pET28-*pilB* | pET-28b derivative for expressing 6His-$PilB_{37\text{-}462}$ | this study |
| pET28-*pilC* | pET-28b derivative for expressing 6His-$PilC_{34\text{-}486}$ | this study |
| pET28-*pilE1* | pET-28b derivative for expressing 6His-$PilE1_{46\text{-}157}$ | this study |
| pET28-*pilE2* | pET-28b derivative for expressing 6His-$PilE2_{46\text{-}150}$ | this study |

**Table S3.** Primers used in this study.

| Name | Sequence |
|---|---|
| **Cloning in pET-28b** | |
| *pilA*-pETF | gggccatggatcatcatcatcatcatcatGATACAGGGCAAAGCCAGAC |
| *pilA*-pETR | cccgtcgacTTACTTCTGTGCCGATCTCAA |
| *pilB*-pETF | gggccatggatcatcatcatcatcatcatAGCAGCCGTGAACTGATTGA |
| *pilB*-pETR | cccggatccTTACGGACCGCTAACAAACC |
| *pilC*-pETF | gggccatggatcatcatcatcatcatcatAATAACATTCTGCGTCAGCGTAGCCA |
| *pilC*-pETR | cccggatccTTAGCTTGCTTTGTATTTATCGC |
| *pilE1*-pETF | ggccatggatcatcatcatcatcatcatCAAGATAACGCTCGTAAGAGCC |
| *pilE1*-pETR | ccggatccTTAGTTTGAGTTTACACCATTAGCAGA |
| *pilE2*-pETF | ggccatggatcatcatcatcatcatcatCAAGATAACGCTCGTAAGAGCC |
| *pilE2*-pETR | ccggatccTTATTTTGAATTAGCACCAGCTTCG |
| **Cloning in pCR8/GW/TOPO** | |
| *pilE1*-F | AGGAACAAAACAAATGCCCCT |
| *pilE1*-R | TCTCAAATGCAGGGTTTTACTACA |
| **Site-directed mutagenesis** | |
| *pilE1*$_{G\text{-}1A}$#1 | GACTTGAAGAAAAAAGGTAAAG**C**TTTTACCTTGGTTGAGTTGATC |
| *pilE1*$_{G\text{-}1A}$#2 | GATCAACTCAACCAAGGTAAAA**G**CTTTACCTTTTTTCTTCAAGTC |
| *pilE1*$_{G\text{-}1S}$#1 | GACTTGAAGAAAAAAGGTAAA**A**GTTTTACCTTGGTTGAGTTGATC |
| *pilE1*$_{G\text{-}1S}$#2 | GATCAACTCAACCAAGGTAAAAC**T**TTTACCTTTTTTCTTCAAGTC |
| *pilE1*$_{E5A}$#1 | GTAAAGGTTTTACCTTGGTTG**C**GTTGATCGTGGTAATTATC |
| *pilE1*$_{E5A}$#2 | GATAATTACCACGATCAAC**G**CAACCAAGGTAAAACCTTTAC |
| **Engineering *S. sanguinis* mutants** | |
| *pheS*-F | ATGACGAAAACGATTGAAGAAC |
| *aph*-R | CTAAAACAATTCATCCAGTAAAA |
| *pilE1*-F1 | CAGGCCGGTGAAAAGACTG |
| *pilE1*-R1 | GTTCTTCAATCGTTTTCGTCATTTTGAATAGATCTCCTGTTTTT |
| *pilE1*-F2 | TTTTACTGGATGAATTGTTTTAGCGACTGGTCTGCTAATGGTG |
| *pilE1*-R2 | GCTCTGTTGAAGGATCCACG |
| *pilE1*-R3 | TTAGTGATGGTGATGGTGATGGTTTGAGTTTACACCATTAGCAG |
| *pilE1*-F3 | CATCACCATCACCATCACTAATTGTCAAATCATCTAAATAAGATGTA |
| *pilE1*-R4 | TTAGTGATGGTGATGGTGATGAGACCAGTCGTAGGTCAAAAC |
| *pilE1*-F4 | CATCACCATCACCATCACTAATTGTCAAATCATCTAAATAAGATGTA |
| *pilE1*-R5 | AGGGGCATTTGTTTTGTTCCT |
| *pilE1*-F5 | TGTAGTAAAACCCTGCATTTGAGA |
| *pilE2*-F1 | GGAACGTCTGACAGGGATGA |
| *pilE2*-R1 | GTTCTTCAATCGTTTTCGTCATATGTATTTTCTCCTAATGTTTTTATG |
| *pilE2*-F2 | TTTTACTGGATGAATTGTTTTAGACCGAAGCTGGTGCTAATTC |
| *pilE2*-R2 | TACCATCCGCAGAAAGACCA |
| *pilE2*-R3 | TTAGTGATGGTGATGGTGATGTTTTGAATTAGCACCAGCTTC |
| *pilE2*-F3 | CATCACCATCACCATCACTAATAACTTGAATTAATTTGAGTTATTCAT |
| *pilE2*-R4 | TTAGTGATGGTGATGGTGATGGGTCCAGTTGTATGTTAAAACG |
| *pilE2*-F4 | CATCACCATCACCATCACTAATAACTTGAATTAATTTGAGTTATTCAT |

Overhangs are in lower case, with restriction sites underlined. Mismatched bases generating point mutations are in bold upper case.
